# Prognostic Biomarkers for Predicting Papillary Thyroid Carcinoma Patients at High Risk Using Nine Genes of Apoptotic Pathway

**DOI:** 10.1101/2020.11.25.397547

**Authors:** Chakit Arora, Dilraj Kaur, G.P.S Raghava

## Abstract

**Objectives:** Aberrant expression of apoptotic genes has been associated with papillary thyroid carcinoma (PTC) in the past, however, their prognostic role and utility as biomarkers remains poorly understood.

**Materials and methods:** In this study, we analysed 505 PTC patients by employing Cox-PH regression techniques, prognostic index models and machine learning methods to elucidate the relationship between overall survival (OS) of PTC patients and 165 apoptosis related genes.

**Results:** It was observed that nine genes (ANXA1, TGFBR3, CLU, PSEN1, TNFRSF12A, GPX4, TIMP3, LEF1, BNIP3L) showed significant association with OS of PTC patients. Five out of nine genes were found to be positively correlated with OS of the patients, while the remaining four genes were negatively correlated. These genes were used for developing risk prediction models. Our voting-based model achieved highest performance (HR=41.59, p=3.36×10^−4^, C=0.84, logrank-p=3.8×10^−8^). The performance of voting-based model improved significantly when we used the age of patients with prognostic biomarker genes and achieved HR=57.04 with p=10^−4^ (C=0.88, logrank-p=1.44×10^−9^). We also developed classification models that can classify high risk patients (survival ≤ 6 years) and low risk patients (survival > 6 years). Our best model achieved AUROC of 0.92. Since these genes can also be used as potential therapeutic targets in PTC, we identified potential drug molecules which could modulate their expression profile.

**Conclusion:** This study briefly revealed the key prognostic biomarker genes in the apoptotic pathway whose altered expression is associated with PTC progression and aggressiveness. In addition to this, risk assessment models proposed here can help in efficient management of PTC patients.

## Introduction

Thyroid cancer’s incidence has been reported to be increasing every year, having the fastest growth rate amongst all the cancers (1). It can be categorized into four major subtypes: i) papillary thyroid carcinoma (PTC), ii) follicular thyroid carcinoma (FTC), iii) medullary thyroid carcinoma (MTC), and iv) anaplastic thyroid carcinoma (ATC). PTC is the most common malignant subtype comprising about 80-85% of all thyroid cancer incidences (2). It is associated with a good prognosis, around 20-30% of the patients are reported to exhibit poor prognosis. This is mostly attributed to the development of distant tumour metastases and recurrences. The progression/transformation of PTC to a more aggressive state, i.e. a poorly differentiated state or a non-differentiated state such as ATC has also been observed in some cases. Thus, efficient risk stratification methods are required for prognostic evaluation and therapeutic decision making in PTC patients. Conventional risk stratifications rely on clinico-pathological factors such as age, gender, tumour size, tumour spread and stage (3,4) but these are plagued with limitations and uncertainties. These limitations demand novel risk assessment methods which are more accurate and derivable from the primary mechanisms driving PTC oncogenesis.

Due to advent of high-throughput sequencing methods and public databases, many biomarkers have been identified for PTC diagnosis, classification, and prognosis prediction. These biomarkers, are important for understanding molecular mechanisms of thyroid cancer. Classic examples include BRAF mutation status, RET/PTC and PAX8/PPAR rearrangements (5–7). BRAF mutations at V599E and T1799A are known to induce the serine kinase levels and thus activate MAPK pathway. Similarly, RET/PTC rearrangement regulate the NFkB activity and thus promote PTC cell migration. Another example of rearrangement is PAX8/PPAR that mediates the transcription pathway and advances PTC progression. In the past, several gene expression-based biomarker have been reported that play a crucial role in PTC prognosis; owing to their altered/differential expression profiles. For example FOXF1 (HR:0.114, 95%CI: 0.045-0.289) and FMO1 (HR:0.202; 95% CI: 0.084-0.487) genes were shown to be associated with favourable RFS (recurrence free survival) in PTC patients (8,9). Downregulation of FOXF1, the gene belonging to the forkhead family of TFs (transcription factors), was also seen to be related with advanced T staging, nodal invasion, and late pathological staging. It has been observed that high expression of FOXE1 a member of forkhead family, also act as a tumour suppressor in PTC (10). High expression of FOXE1 was found to negatively regulate PDFGA (target gene platelet-derived growth factor A) expression in the early stage of PTC and thus affect the migration, proliferation and invasion of PTC. Proteoglycans genes (e.g., SDC1, SDC4, KLK7, KLK10, SLPI, GDF15) were found to be overexpressed in PTC samples (11). Similarly, lower expression of VHL gene was shown to be associated with aggressive PTC features and DFI (disease free interval, logrank-p=0.007) (12). VHL (von Hippel–Lindau) protein, by acting as a substrate recognition unit in a multiprotein complex with E3 ubiquitin ligase activity, is involved in the degradation of the proteins such as HIF-α. Whereas HIF-α regulates the levels of various angiogenic factors and is thus negatively affected, resulting in a reduced angiogenesis. Bhalla et al (13) reported 36 RNA transcripts whose expression profiles were used to distinguish early and late-stage PTC patients (AUROC 0.73). In addition to above, number pf candidate genes and biomarkers have been reported in previous studies (14–16). Despite tremendous progress in the field of prognostic biomarkers still it is far from perfection. There is a need to develop methods to identify genes of critical pathways that can serve as prognostic biomarkers, particularly genes involved in apoptosis.

Apoptosis or programmed cell death is the process for eliminating cells in multicellular organisms. Dysregulation of apoptosis is responsible for many diseases including cancer. Numerous studies have identified key biomarkers linked with the cellular apoptosis. Charles EM *et al* present the literature related to the apoptotic molecules implicated as biomarkers in melanoma (17). Another review provides extensive information related to apoptotic biomarkers such as p53, Bcl2, Fas/FasL, TRAIL in colorectal cancer (18). Several other studies have also identified key molecules with prognostic roles in other cancers like gastric cancer (19,20), breast cancer (21), lung cancer (22), bladder urothelial carcinoma (23), glioblastoma (24) and osteosarcoma (25). Apoptosis has also been found to have a crucial role in carcinogenesis of thyroid cancer. Alterations in an increasing number of apoptotic molecules such as p53, Bcl2, Bcl-XL, Bax, p73, Fas/FasL, PPARG, TGFb and NFKb have been associated with thyroid cancer (26). Since apoptotic resistance is mostly accounted for tumour proliferation and aggressiveness, apoptotic pathway has also emerged as a crucial target to develop anticancer treatments for thyroid tumours. For example, paclitaxel and manumycin are known to stimulate *p21* expression and induce apoptosis in ATC (27). Lovastin inhibits protein geranylation of the Rho family and thus induces apoptosis in ATC (28). UCN-01 inhibits expression of Bcl-2, leading to apoptosis (29). Since apoptosis in PTC is a complicated multistep process involving a number of genes, it remains poorly understood and needs to be further explored at a genetic level.

In this study, we exploited the mRNA expression data obtained from The Cancer Genome Atlas-Thyroid Carcinoma (TCGA-THCA) cohort and identified key apoptotic genes that are associated with PTC prognosis. We further constructed multiple risk stratification models using these genes and evaluated the potential of these models for prognosis using univariate and multivariate analyses, Kaplan Meier survival curves and other standard statistical tests. The 9 gene voting based model was found to perform the best and also stratified high risk clinical groups significantly. Finally, after a comprehensive prognostic comparison with other clinico-pathological factors, we developed a hybrid model which combines expression profile of nine genes with ‘Age’ to predict High and Low risk PTC patients with high precision. We also catalogued candidate small molecules that can modulate the expression of these genes and could be potentially employed in efficient treatment of PTC patients.

## 2 Materials and Methods

### 2.1 Dataset and pre-processing

The original dataset consisted of quantile normalized RNAseq expression values for 573 Thyroid Carcinoma (THCA) patients that were obtained from ‘The Cancer Genome Atlas’ (TCGA) using TCGA Assembler-2 (30). Out of which, information about overall survival (OS) time and censoring was available for 505 patients. The list of genes involved in the apoptotic pathway were taken from previous study (31). Thus, the final dataset was reduced to 505 samples, using in-house python and R-scripts, constituting RNAseq values for 165 apoptotic genes. More details about clinical, pathological and demographic features corresponding to the final dataset are summarized in Supplementary S1 Table 1.

### 2.2 Survival analysis

Hazard ratios (HR) and confidence intervals (95% CI) were evaluated to predict the risk of death related with high- and low-risk groups based on overall survival time of patients. These were stratified on the basis of appropriate cut-offs for various factors, using the univariate unadjusted Cox-Proportional Hazard (Cox-PH) regression models. Kaplan-Meier (KM) plots were used to compare survival curves of the risk groups. ‘survival’ and ‘survminer’ packages were used to perform survival analyses on the dataset. log-rank tests were used to estimate the statistical significance between the survival curves. Concordance index (C) was computed to measure the strength of predictive ability of the model (32–34); p-values less than 0.05 were considered as significant. Multivariate survival analysis based on cox regression was employed to compare the relationship between various covariates.

### 2.3 Multiple gene-based models

#### 2.3.1 Machine learning based regression (MLR) models

Various regression models from ‘sklearn package in Python (35) were implemented to fit the gene expression values against the OS time. Regressors such as Linear, Ridge, Lasso, Lasso-Lars, Elastic-Net, Random-forest (RF) and K-nearest neighbours (KNN) were used. Five-fold cross-validation was used for training and validation studies, as done in previous studies (36–40). All five test datasets were combined as ‘predicted OS’ and stratification was performed using it. Median cutoff was used to estimate HR, CI and p-values. Hyperparameter optimization and regularization was achieved using the in-built function ‘GridsearchCV’. Model’s performance is denoted using standard parameters viz. RMSE (root mean squared error) and MAE (mean absolute error).

#### 2.3.2 Prognostic Index (PI)

For *n* genes, Prognostic Index (PI) is defined as:

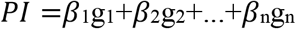

Where g_i_ represent genes and *β_i_* represent regression coefficients obtained from Cox univariate regression analysis as done in (41,42). Risk groups were stratified based on best PI cutoff estimated using cutp from ‘survMisc’ package in R. HR, p-values, C index were then evaluated using this cutoff.

#### 2.3.3 Gene voting based model

Corresponding to an individual gene expression (median cutoff), a risk label ‘High Risk’ or ‘Low Risk’ was assigned to each patient. Thus, for *n* survival associated genes, every patient was denoted by a ‘risk’ vector of *n* risk labels. In gene voting based method, the patient is ultimately classified into one of the high/low risk categories based on the dominant ‘label’ (i.e. occurring more than at least n/2 times) in this vector. This is followed by evaluation of standard metrics.

## 3. Results

### 3.1 Survival associated apoptotic genes

Cox-Proportional Hazard models were used to find those apoptotic pathway genes that are related with PTC patient survival (Supplementary S1 Table 2). A univariate Cox-PH analysis revealed a total of 5 good prognostic marker (GPM) genes i.e the genes that are positively correlated with patient OS time and 4 bad prognostic marker (BPM) genes which are negatively correlated with OS time of the patients. GPM genes are ANXA1, CLU, PSEN1, TNFRSF12A and GPX4 while BPM genes are TGFBR3, TIMP3, LEF1 and BNIP3L. Table 1 shows the results for these genes along with the metrics associated with stratification of high/low risk patients at median cutoff. We compared the expression of these genes in normal patients (TCGA and GTEX normal samples) with cancer patients, with the help of GEPIA server (43). Out of these 9 genes, 7 genes (except BNIP3L and LEF1) were found to be differentially expressed significantly, thus elucidating their role in PTC oncogenesis. Results are shown in Supplementary S2 Figure 1. The precise molecular information about these 9 genes and PMIDs of the studies pertaining to their role in cancer, as obtained from GeneCards (44) and The Candidate Cancer Gene Database (CCGD) (45) respectively, is provided in Supplementary S1 Table 3. Supplementary S1 Table 4 shows results of risk stratification performed using various previously suggested prognostic genes in PTC using cox univariate analysis in TCGA-THCA dataset at median expression cutoff for overall-survival.

**Table 1.** The table shows results of univariate cox regression with >median cutoff. Genes with HR>1 are BPM while HR<1 are GPM.

|  | Gene | HR | p-val | C | %95 CI L | %95 CI U | Logrank-p |
| --- | --- | --- | --- | --- | --- | --- | --- |
| 1. | ANXA1 | 0.14 | $2.82 \times 10^{-3}$ | 0.72 | 0.04 | 0.51 | $7.35 \times 10^{-4}$ |
| 2. | TGFBR3 | 5.68 | $7.90 \times 10^{-3}$ | 0.62 | 1.58 | 20.49 | $2.82 \times 10^{-3}$ |
| 3. | CLU | 0.18 | $8.15 \times 10^{-3}$ | 0.53 | 0.05 | 0.64 | $2.92 \times 10^{-3}$ |
| 4. | PSEN1 | 0.15 | $1.20 \times 10^{-2}$ | 0.71 | 0.03 | 0.66 | $2.38 \times 10^{-3}$ |
| 5. | TNFRSF12A | 0.25 | $1.57 \times 10^{-2}$ | 0.51 | 0.08 | 0.77 | $1.30 \times 10^{-2}$ |
| 6. | GPX4 | 0.27 | $2.98 \times 10^{-2}$ | 0.62 | 0.09 | 0.88 | $2.09 \times 10^{-2}$ |
| 7. | TIMP3 | 3.49 | $3.52 \times 10^{-2}$ | 0.68 | 1.09 | 11.18 | $2.53 \times 10^{-2}$ |
| 8. | LEF1 | 3.36 | $4.10 \times 10^{-2}$ | 0.68 | 1.05 | 10.77 | $3.00 \times 10^{-2}$ |
| 9. | BNIP3L | 4.56 | $4.78 \times 10^{-2}$ | 0.68 | 1.01 | 20.46 | $2.05 \times 10^{-2}$ |

### 3.2 Risk estimation using multiple gene-based models

Several risk stratification models based on MLR, prognostic index and gene voting were constructed using the expression profile of nine survival associated apoptotic genes. Table 2 shows the results corresponding to various risk models. Amongst these, the performance of gene voting based model was found to be the best with HR=41.59 and p~10^−4^ with C-value of 0.84. In addition, high/low risk groups survival curves were significantly separated with a logrank-p~10^−8^ using voting based model. As shown in KM plot (Figure 1), 10-year survival rate for low risk patients was close to 98%, for high risk patients it was drop to 40%. PI based model performed the second best with HR=17.55 and p~10^−3^ (Supplementary S2 Figure 2), and regression-based RF model was the third best (and top amongst MLR models) with HR=3.09 but p-value was found to be statistically insignificant.

**Table 2.** The performance of different models developed using multiple gene expression profile-based.

|  | Model | HR | p-val | C | %95 CI L | %95 CI U | Logrank-p |
| --- | --- | --- | --- | --- | --- | --- | --- |
| 1. | **Voting based** | **41.59** | **$3.36 \times 10^{-4}$** | **0.84** | **5.42** | **319.17** | **$3.80 \times 10^{-8}$** |
| 2. | PI | 17.55 | $5.88 \times 10^{-3}$ | 0.65 | 2.29 | 134.72 | $6.73 \times 10^{-5}$ |
| 3. | RF | 3.09 | $8.43 \times 10^{-2}$ | 0.68 | 0.86 | 11.09 | $5.91 \times 10^{-2}$ |
| 4. | Linear | 1.59 | $3.98 \times 10^{-1}$ | 0.54 | 0.54 | 4.65 | $4.04 \times 10^{-1}$ |
| 5. | KNN | 1.09 | $8.68 \times 10^{-1}$ | 0.56 | 0.38 | 3.12 | $8.68 \times 10^{-1}$ |
| 6. | Lasso | 1.07 | $9.06 \times 10^{-1}$ | 0.52 | 0.37 | 3.08 | $9.06 \times 10^{-1}$ |
| 7. | ElasticNet | 1.07 | $9.06 \times 10^{-1}$ | 0.52 | 0.37 | 3.08 | $9.06 \times 10^{-1}$ |
| 8. | LassoLars | 1.06 | $9.18 \times 10^{-1}$ | 0.52 | 0.37 | 3.06 | $9.18 \times 10^{-1}$ |
| 9. | Ridge | 0.84 | $7.43 \times 10^{-1}$ | 0.50 | 0.29 | 2.42 | $7.44 \times 10^{-1}$ |

\*boldface represents statistically significant results (p-val, logrank $p < 0.05$ ). MLR hyperparameters and evaluation statistics are provided in S1 Table 5.

**Figure 1.**
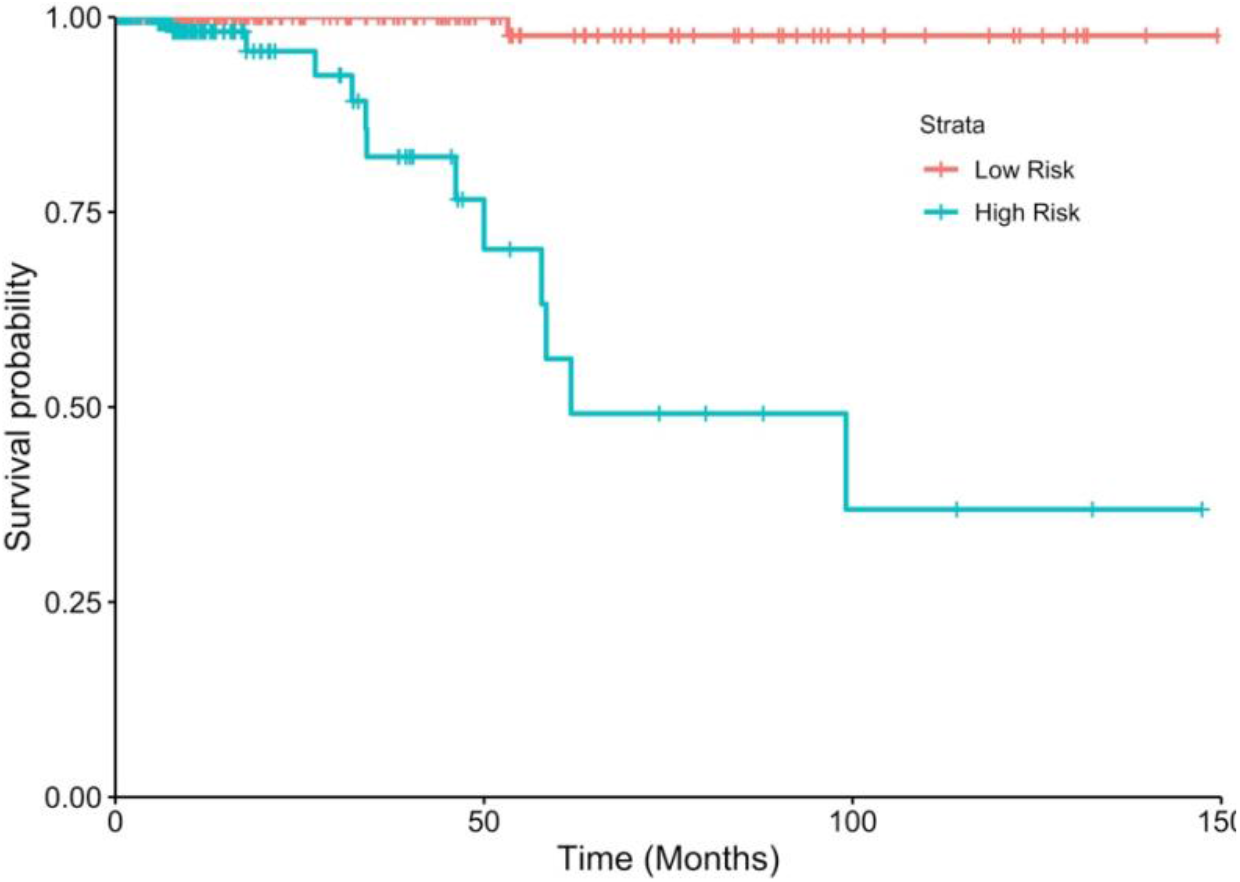
KM plot showing risk stratification of PTC patients based on gene voting model. Patients with greater than five ‘high risk’ labels in the 9-bit risk vector are assigned (blue) as High Risk (HR=41.59, p=3.36×10^−4^, C=0.84, logrank-p=3.8×10^−8^) while others were assigned as Low Risk (red).

### 3.3 Multiple gene model sub-stratifies patients in clinico-pathological high-risk groups

Past studies indicate the role of certain clinico-pathological factors in PTC prognosis such as age, gender, ethnicity and tumour size (3,4). Thus, we performed a univariate analysis to assess the association of these factors with OS in our dataset. Table 3 shows the results of the univariate analysis. Patient age is seen to be the most significant factor in the PTC prognosis (HR=48.65, C=0.86), and is supported by numerous earlier studies (46). The AJCC thyroid cancer staging also includes an age cutoff of 55 years to classify tumour stages (47), since patients with age<55y usually show a very good prognosis. However, we obtained the best stratification at the age cutoff of 60y which also corroborated with a recent study (48). AJCC Tumour staging was seen to be the second-best risk predictor with HR=9.23 and C=0.76.

In order to evaluate the strength of the 9-gene based model, we sub-stratified the patients in the clinical high-risk subgroups i.e Age>60 and Stage III/IV patients. Figure 2 shows the sub-stratification by means of KM plots. A significant separation between the survival curves is seen, as denoted by logrank test’s p-values. KM plots for other high-risk subgroups are provided in Supplementary S2 Figure 3.

**Table 3.**
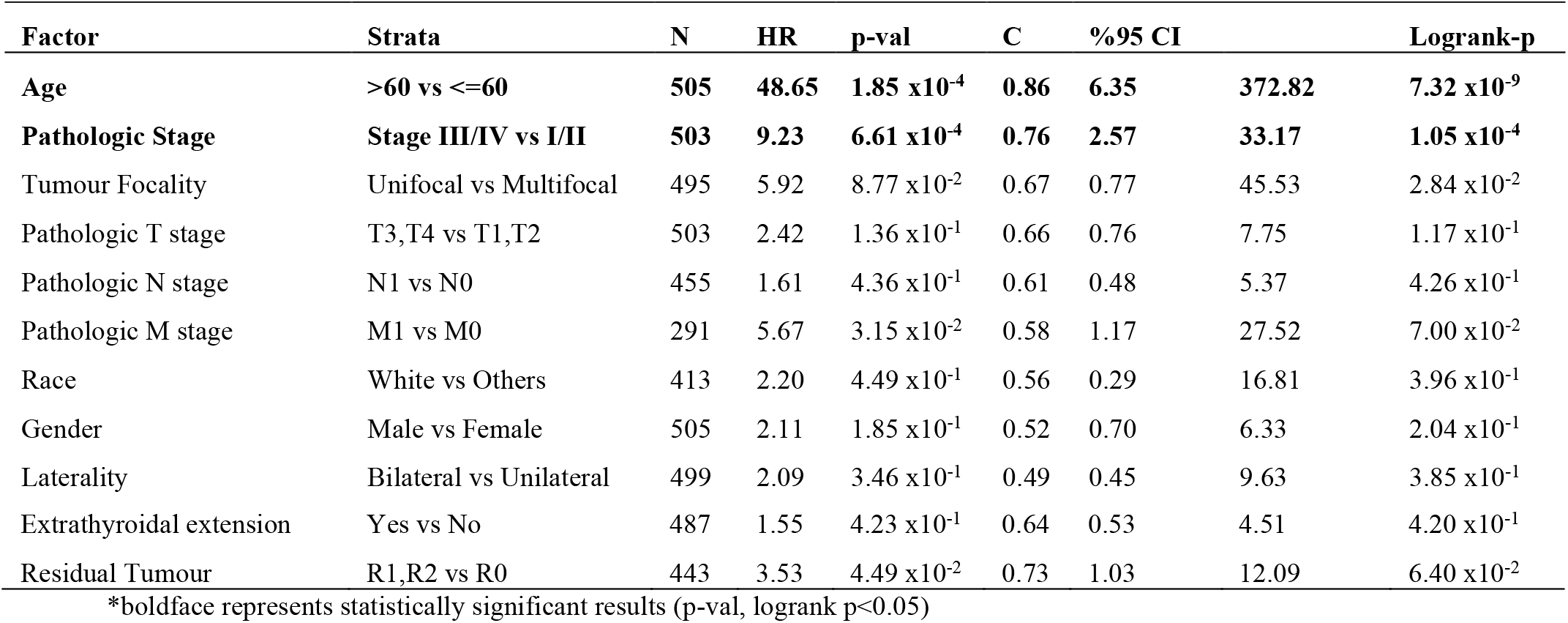
Univariate analysis using clinico-pathological features. Age is seen to be the most significant factor. In laterality, unilateral: right lobe, left lobe and isthmus.

**Figure 2.**
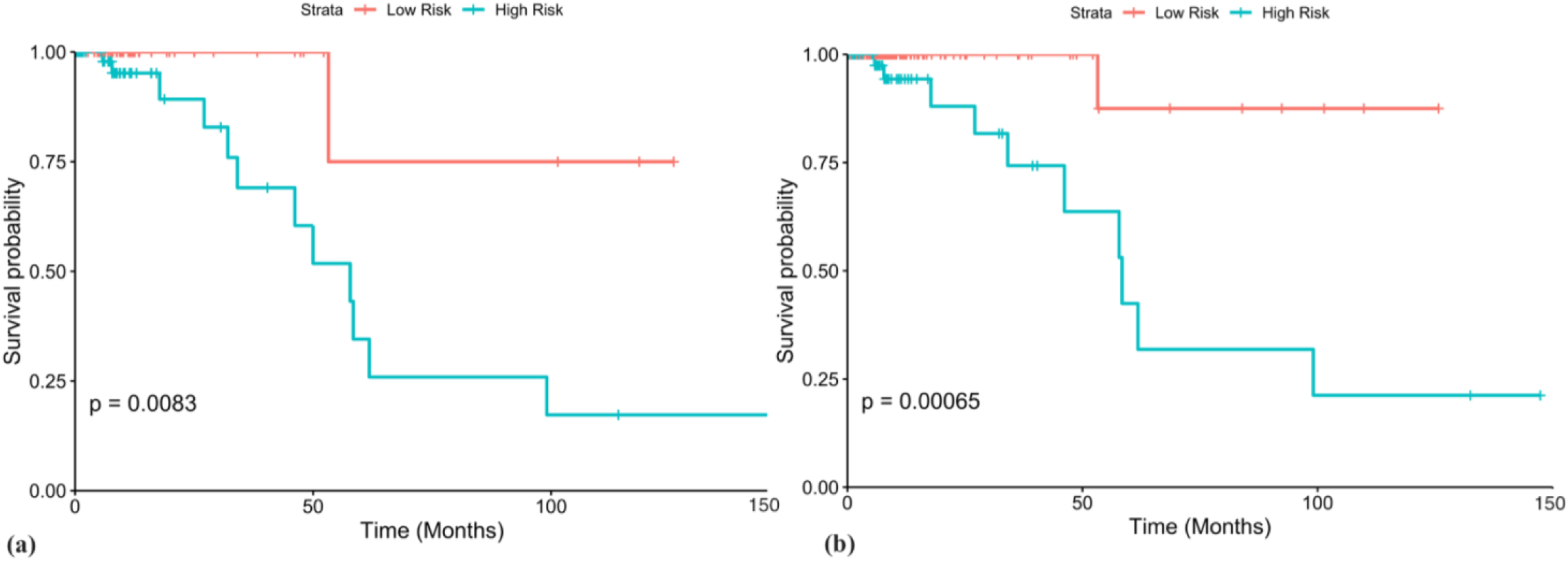
Voting model sub-stratifies high risk groups. (a) Patients with with age>60y (n=113) were stratified into high and low risk groups with HR=9.49, p=3.08×10^−2^ and C=0.72. (b) Stage III/IV patients (n=167) were stratified into high and low risk groups with HR=15, p=0.01 and C=0.81. p-values from logrank tests are shown in the KM plots.

### 3.4 Hybrid voting model

After obtaining three prominent prognostic markers i.e. multiple gene voting model, patient age and AJCC stage, we performed a multivariate cox regression survival analysis. The analysis showed that patient age (HR=13.3, p=0.02) and gene voting model (HR=13.3, p=0.015) were independent covariates, while p-value corresponding to staging was insignificant as depicted by the forest plot in Figure 3(a). Next, we developed a hybrid voting model by combining patient age with the 9-gene voting model for risk stratification purposes. The risk vector associated with each patient was thus now a 10-bit vector with 1 bit assigned to risk label due to age. Supplementary 1 Table 6 shows results pertaining to stratification by hybrid models with different age cutoffs (45y-65y). We observed that the model performed best when the age cutoff was set at 65y (HR=57.04, C=0.88) as compared to 60y (HR=54.82, C=0.87). While the risk groups have a better separation in the former model, the 5 and 10-year survival is comparable in both models. High risk groups show a 40% 5-year survival and around 25% 10-year survival, whereas, low risk groups have a 98% 5 and 10-year survival chance. Figure 3 shows the KM plots corresponding to both the hybrid models.

**Figure 3.**
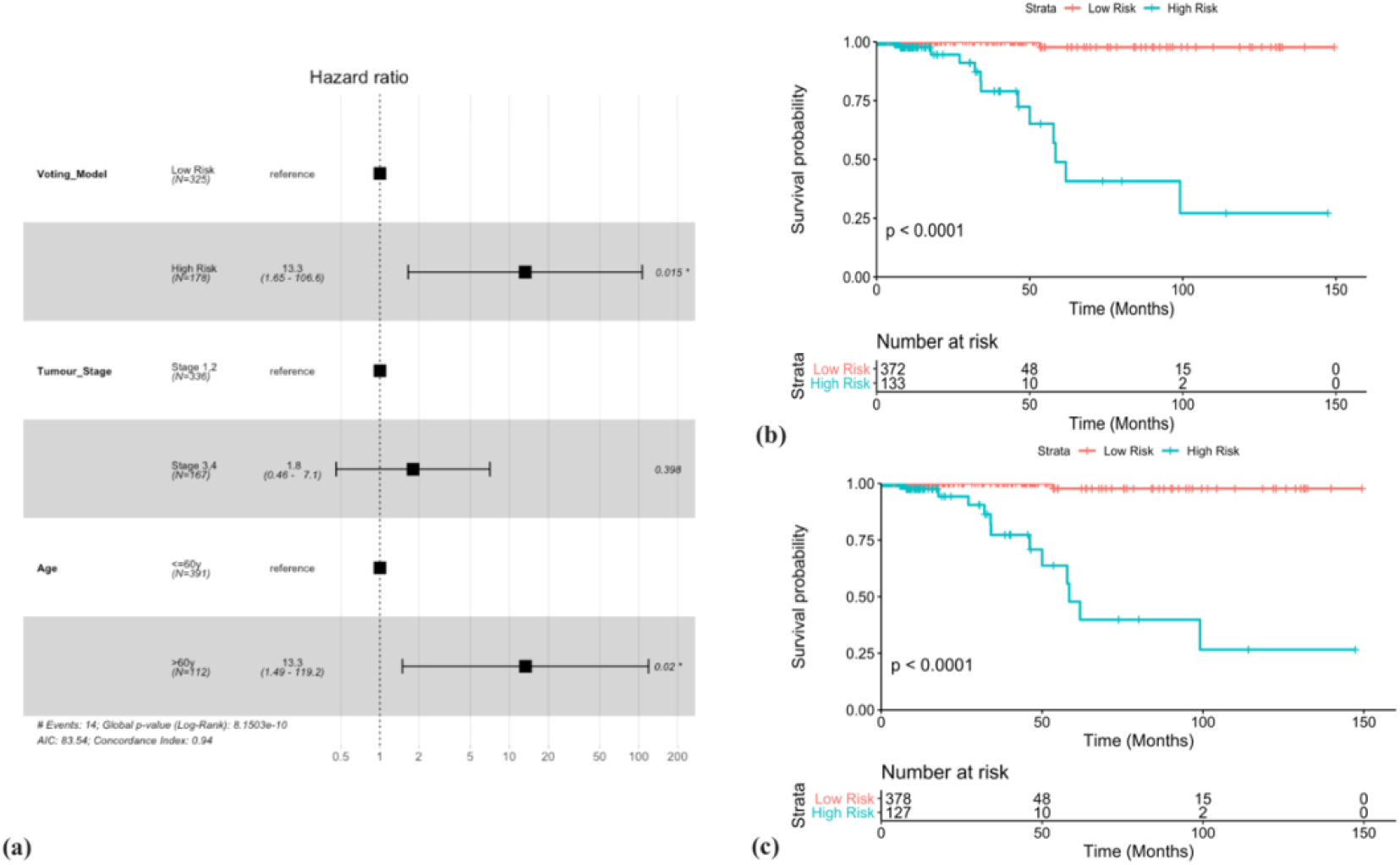
Hybrid models for risk stratification. (a) Multivariate analysis reveals Age (HR=13.3, p=0.02) and Voting model (HR=13.3, p=0.015) as two independent covariates, while tumour stage was found to be insignificant. (b) Risk stratification by hybrid model with age cutoff >60y (HR=54.82, p=1.18×10^−4^, C=0.87, %95CI: 7.14-420.90 and logrank-p=2.3×10^−9^). (b) (b) Risk stratification by hybrid model with age cutoff >65y (HR=57.04, p~10^−4^, C=0.88, %95CI: 7.44-437.41 and logrank-p=1.4×10^−9^)

### 3.5 Predictive validation

As implemented in (49) we performed a predictive assessment of our models using sub-samples of the complete dataset. Sampling sizes of 50%, 70% and 90% were chosen with 100 iterations each. HR and C index were evaluated for each iteration corresponding to the 9-gene voting model and hybrid models. Figure 4 shows the boxplots corresponding to the results. It is evident from the figure that the hybrid model with age cutoff of >65 years performs the best as compared to other models in terms of HR and C values. The median HR (27.03, 39.53, 50.33) and C (0.86, 0.87, 0.87) values for this model remain better than the other two models’ despite of the sampling size. Thismethod ensured that the risk stratification models were robust and performed well with random datasets of different sizes.

**Figure 4.**
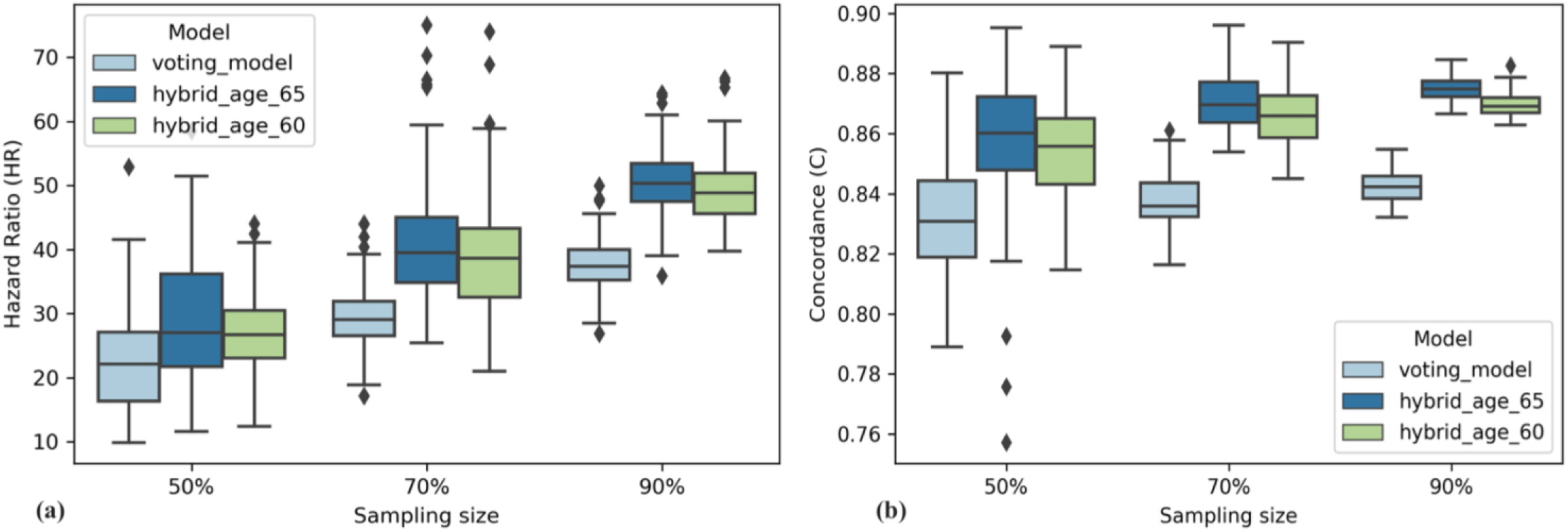
Predictive validation of voting based model and hybrid models. (a) Grouped boxplots corresponding to estimated Hazard Ratio (y-axis) for 100 iterations of data sampling (x-axis). (b) Similarly, estimation of Concordance index (y-axis) for different models using random sampling (x-axis).

### 3.6 Classification using hybrid model

In order to evaluate the classification performance of above hybrid feature, we developed classification models. Firstly, we segregated patients into poor survival (negative data) and good survival (positive data) using an OS time cut-off. Secondly, we used package ‘survivalROC’ to calculate the true positive (TPR) and true negative rates (TNR). Here, a true positive prediction being the patient whose OS> cut-off time as well as who was in low-risk group according to hybrid model, while converse applies for a true negative prediction. Consequently, an AUROC value (Area under receiver operating characteristic curve) was calculated, which denoted the model’s classification ability. Out of various time cut-offs used (2-10 years), the model was seen to perform best at the cut-off of 6 years. At this cut-off, a maximum AUROC value of 0.92 was obtained. The ROC curve is represented in Figure 5(b).

**Figure 5.**
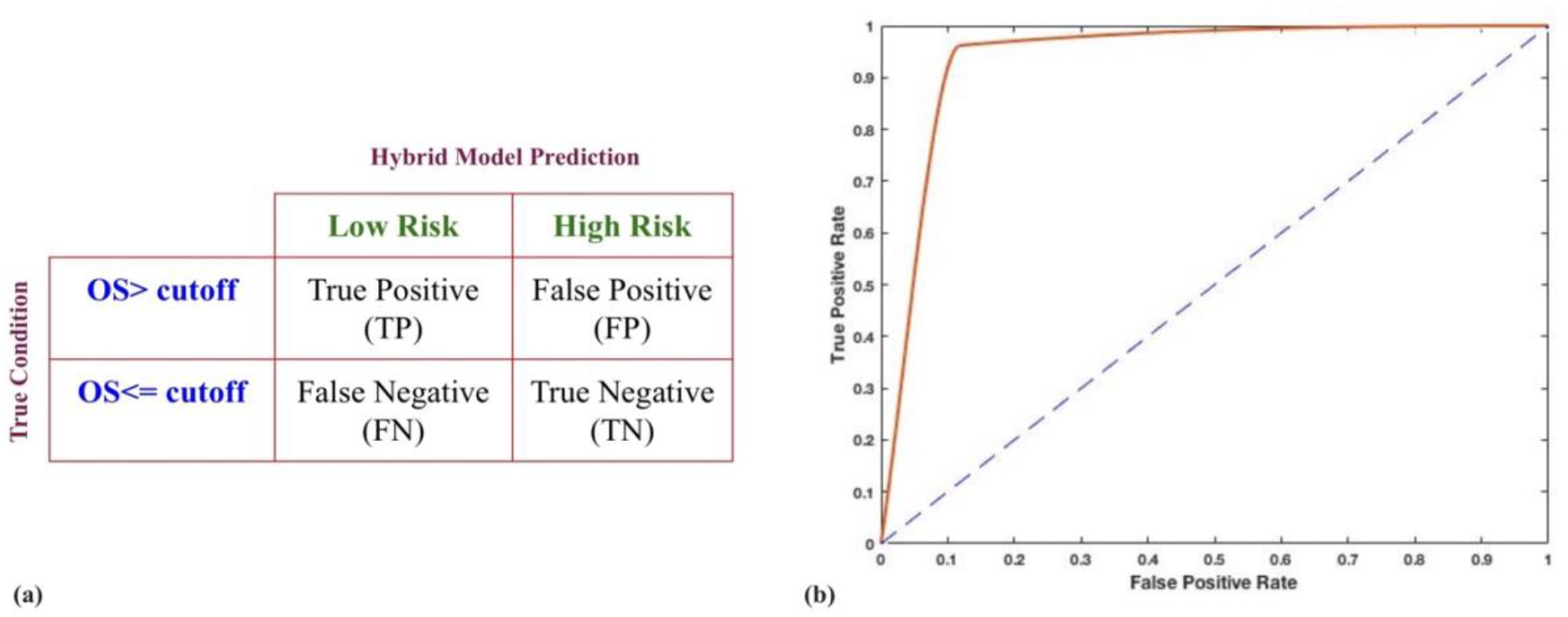
Hybrid models for classification of PTC patients using OS. (a) Terminology used for evaluation of confusion matrix. Initial risk labelling was done using an OS cutoff with patients having OS> cutoff labelled as positive or low risk and vice-versa for patients with OS≤cutoff. (b) ROC curve for hybrid model using asge cutoff og 65y. AUROC of 0.92 was obtained.

### 3.7 Screening of therapeutic drug molecules

Another major step after the identification of key genes whose altered expression is associated with PTC risk is the choice of therapy which can alter this situation. This requires selection of small molecules which can induce or inhibit the gene expression of downregulated and upregulated genes in PTC. As implemented in (50), we found drug molecules which could reverse gene expression induced by PTC using the ‘Cmap2 database’ (51,52). A list of probe ids corresponding to upregulated genes (TGFBR3, TIMP3, LEF1 and BNIP3L) and downregulated genes (ANXA1, CLU, PSEN1, TNFRSF12A and GPX4) was used as input to fetch small molecules ranked on the basis of p-values (results in Supplementary S1 Table 7). Top 2 negative and positively enriched molecules were Lomustine (enrichment=-0.908, p=0.0001) and Deferoxamine (enrichment= 0.663, p=0.0006). Lomustine is an alkylating nitrosourea compound which is already used in chemotherapy, especially in brain tumours, and has been associated with inducing apoptosis in past studies (53). Deferoxamine (DFO) is an iron chelator which reduces iron content in cells. Various studies have confirmed that diminishing iron content inhibits tumor cell proliferation and induces apoptosis (54,55). Out of the various iron-chelators available, DFO is the most widely used iron-chelator and has shown to display these anti-tumor effects (56,57).

## 4. Discussion

Though, PTC is known to have a very good prognosis, there still remains a decent proportion of patients with an exceptionally poor prognosis. As a result of which, accurate risk assessment strategies are required for clinical decision making and therapeutic intervention. While conventional clinico-pathological factors such as age, stage, extrathyroidal spread and tumour size are significant in risk stratification of PTC patients, they have their own limitations and are not that efficient. Thus, aided by the development in the high-throughput sequencing methods and availability of a huge amount of experimental data, various molecular prognostic markers have been proposed in the past (5–12,14–16). The understanding of the mechanistic roles of these molecules in the PTC carcinogenesis has initiated a further enquiry into other complicated molecular processes, which may be crucial in PTC progression and development. As uncovered in the past investigations, apoptosis in PTC is a multifaceted and multistep process. Apoptosis based biomarkers have also been proposed for many other cancers (17–19,21,22). Despite the fact that the role of genes and their associated proteins such as Fas/FasL, Bcl-2 family, p53, and others have been exhibited in PTC malignant growth, our comprehension of the collaboration between these molecules is still poor. The crosstalk that happens between numerous upstream signals and downstream effectors presents an extensive challenge to the ongoing investigation of apoptosis in PTC. Be that as it may, these complexities provide opportunities for the disclosure of novel prognostic biomarkers and therapeutic targets.

In the current study, we examined the genes involved in the apoptotic pathway and evaluated the prognostic potential of the expression of these genes in PTC. We employed a recent gene expression dataset, and found out that out of 165 genes, 9 genes were significantly associated with PTC prognosis. Out of these genes, ANXA1 or annexin A1 expression has been shown to be associated with differentiation in PTC (58). Western blotting experiments showed high levels of ANXA1 in papillary thyroid carcinoma and follicular cells while undifferentiated thyroid carcinoma cells had low levels of ANXA1 protein. TGFBR3 gene was found to be differentially expressed between normal and PTC samples and was shown to be related with progression free interval (15). The encoded TGFBR3 protein is a membrane proteoglycan and is known to function as a co-receptor along-with other TGF-beta receptor superfamily members. Reduced expression of the TGFBR3 protein has also been observed in various other cancers. CLU protein is a secreted chaperone which has been previously suggested to be involved in apoptosis and tumour progression. Altered CLU expression has also been proposed as biomarker for assessment of indeterminate thyroid nodules (59). PSEN1 mutations have been shown to be linked with MTC (60). TNFRSF12A was linked to aging and thyroid cancer (61) and also shown to be a PTC prognostic biomarker in yet another study (62). GPX4 is an essential seleno-protein shown to be associated with aging and cancer (63). TIMP3 levels were found to be associated with BRAF mutations in PTC (64). LEF1 expression was found to be up-regulated in PTC (65) and BNIP3L-CDH6 interaction has been linked with defunct autophagy and epithelial to mesenchymal transition (EMT) in PTC (66). We also evaluated the risk stratification performance of other genes suggested in past studies and show that the 9 genes proposed in our study show better results. We also found potential drug-molecules which could be potentially used for PTC therapy and require future efforts. Lomustine and Deferoxamine were two such top molecules which are widely used in anti-cancer treatment due to their apoptosis inducing roles. Further, a multiple gene expression profile-based voting model was developed for these 9 genes. Apart from its superior performance in the complete dataset, this model was able to segregate high and low risk patients in clinically established high risk groups. We further gauged the performance of this multiple gene model against clinico-pathological factors, using a multivariate survival analysis. The analysis led to identification of ‘Patient Age’ as another independent significant factor, and thus a hybrid model utilizing the 9 gene expression profile and age was developed. This model further boosted the performance and provided better stratification. Further, Monte Carlo validation was performed to assess the robustness of this model. The model was also able to achieve an AUROC of 0.92 for classification of patients having more than 6 years overall survival with those having less than or equal to 6 years overall survival time. In conclusion, we identified key genes with a possible role in PTC pathogenesis and prognosis. While, this is supported by previous literature and explored in the current study as an in-silico analysis, it is subjected to further validation. Also, apart from their strong prognostic potential, as elucidated in this study, these genes could also be investigated further in the context of therapeutic targets in PTC and clinical decision making.

## References

1. Mao Y, Xing M. Recent incidences and differential trends of thyroid cancer in the USA. Endocr Relat Cancer (2016) 23:313–322. doi:10.1530/ERC-15-0445

2. LiVolsi VA. Papillary thyroid carcinoma: an update. Mod Pathol (2011) 24 Suppl 2:S1–9. doi:10.1038/modpathol.2010.129

3. Carrillo JF, Frias-Mendivil M, Ochoa-Carrillo FJ, Ibarra M. Accuracy of fine-needle aspiration biopsy of the thyroid combined with an evaluation of clinical and radiologic factors. Otolaryngol Head Neck Surg (2000) 122:917–921. doi:10.1016/s0194-5998(00)70025-8

4. Are C, Shaha AR. Anaplastic thyroid carcinoma: biology, pathogenesis, prognostic factors, and treatment approaches. Ann Surg Oncol (2006) 13:453–464. doi:10.1245/ASO.2006.05.042

5. Cohen Y, Xing M, Mambo E, Guo Z, Wu G, Trink B, Beller U, Westra WH, Ladenson PW, Sidransky D. BRAF mutation in papillary thyroid carcinoma. J Natl Cancer Inst (2003) 95:625–627. doi:10.1093/jnci/95.8.625

6. Soares P, Trovisco V, Rocha AS, Lima J, Castro P, Preto A, Maximo V, Botelho T, Seruca R, Sobrinho-Simoes M. BRAF mutations and RET/PTC rearrangements are alternative events in the etiopathogenesis of PTC. Oncogene (2003) 22:4578–4580. doi:10.1038/sj.onc.1206706

7. Fukushima T, Suzuki S, Mashiko M, Ohtake T, Endo Y, Takebayashi Y, Sekikawa K, Hagiwara K, Takenoshita S. BRAF mutations in papillary carcinomas of the thyroid. Oncogene (2003) 22:6455–6457. doi:10.1038/sj.onc.1206739

8. Gu Y, Hu C. Bioinformatic analysis of the prognostic value and potential regulatory network of FOXF1 in papillary thyroid cancer. Biofactors (2019) 45:902–911. doi:10.1002/biof.1561

9. Luo J, Zhang B, Cui L, Liu T, Gu Y. FMO1 gene expression independently predicts favorable recurrence-free survival of classical papillary thyroid cancer. Future Oncol (2019) 15:1303–1311. doi:10.2217/fon-2018-0885

10. Ding Z, Ke R, Zhang Y, Fan Y, Fan J. FOXE1 inhibits cell proliferation, migration and invasion of papillary thyroid cancer by regulating PDGFA. Mol Cell Endocrinol (2019) 493:110420. doi:10.1016/j.mce.2019.03.010

11. Reyes I, Reyes N, Suriano R, Iacob C, Suslina N, Policastro A, Moscatello A, Schantz S, Tiwari RK, Geliebter J. Gene expression profiling identifies potential molecular markers of papillary thyroid carcinoma. Cancer Biomark (2019) 24:71–83. doi:10.3233/CBM-181758

12. Todorovic L, Stanojevic B, Mandusic V, Petrovic N, Zivaljevic V, Paunovic I, Diklic A, Saenko V, Yamashita S. Expression of VHL tumor suppressor mRNA and miR-92a in papillary thyroid carcinoma and their correlation with clinical and pathological parameters. Med Oncol (2018) 35:17. doi:10.1007/s12032-017-1066-3

13. Bhalla S, Kaur H, Kaur R, Sharma S, Raghava GPS. Expression based biomarkers and models to classify early and late-stage samples of Papillary Thyroid Carcinoma. PLoS One (2020) 15:e0231629. Available at: https://doi.org/10.1371/journal.pone.0231629

14. Soares P, Celestino R, Melo M, Fonseca E, Sobrinho-Simoes M. Prognostic biomarkers in thyroid cancer. Virchows Arch (2014) 464:333–346. doi:10.1007/s00428-013-1521-2

15. Wu M, Yuan H, Li X, Liao Q, Liu Z. Identification of a Five-Gene Signature and Establishment of a Prognostic Nomogram to Predict Progression-Free Interval of Papillary Thyroid Carcinoma. Front Endocrinol (Lausanne) (2019) 10:790. doi:10.3389/fendo.2019.00790

16. Li X, He J, Zhou M, Cao Y, Jin Y, Zou Q. Identification and Validation of Core Genes Involved in the Development of Papillary Thyroid Carcinoma via Bioinformatics Analysis. Int J Genomics (2019) 2019:5894926. doi:10.1155/2019/5894926

17. Charles EM, Rehm M. Key regulators of apoptosis execution as biomarker candidates in melanoma. Mol Cell Oncol (2014) 1:e964037. doi:10.4161/23723548.2014.964037

18. Zeestraten ECM, Benard A, Reimers MS, Schouten PC, Liefers GJ, van de Velde CJH, Kuppen PJK. The prognostic value of the apoptosis pathway in colorectal cancer: a review of the literature on biomarkers identified by immunohistochemistry. Biomark Cancer (2013) 5:13–29. doi:10.4137/BIC.S11475

19. Bai Z, Ye Y, Liang B, Xu F, Zhang H, Zhang Y, Peng J, Shen D, Cui Z, Zhang Z, et al. Proteomics-based identification of a group of apoptosis-related proteins and biomarkers in gastric cancer. Int J Oncol (2011) 38:375–383. doi:10.3892/ijo.2010.873

20. Ding L, Li B, Yu X, Li Z, Li X, Dang S, Lv Q, Wei J, Sun H, Chen H, et al. KIF15 facilitates gastric cancer via enhancing proliferation, inhibiting apoptosis, and predict poor prognosis. Cancer Cell Int (2020) 20:125. doi:10.1186/s12935-020-01199-7

21. Pandya V, Githaka JM, Patel N, Veldhoen R, Hugh J, Damaraju S, McMullen T, Mackey J, Goping IS. BIK drives an aggressive breast cancer phenotype through sublethal apoptosis and predicts poor prognosis of ER-positive breast cancer. Cell Death Dis (2020) 11:448. doi:10.1038/s41419-020-2654-2

22. Nakano T, Go T, Nakashima N, Liu D, Yokomise H. Overexpression of Antiapoptotic MCL-1 Predicts Worse Overall Survival of Patients With Non-small Cell Lung Cancer. Anticancer Res (2020) 40:1007–1014. doi:10.21873/anticanres.14035

23. Zeng S, Liu A, Dai L, Yu X, Zhang Z, Xiong Q, Yang J, Liu F, Xu J, Xue Y, et al. Prognostic value of TOP2A in bladder urothelial carcinoma and potential molecular mechanisms. BMC Cancer (2019) 19:604. doi:10.1186/s12885-019-5814-y

24. Liu Y-Q, Wu F, Li J-J, Li Y-F, Liu X, Wang Z, Chai R-C. Gene Expression Profiling Stratifies IDH-Wildtype Glioblastoma With Distinct Prognoses. Front Oncol (2019) 9:1433. doi:10.3389/fonc.2019.01433

25. Ma L, Zhang L, Guo A, Liu LC, Yu F, Diao N, Xu C, Wang D. Overexpression of FER1L4 promotes the apoptosis and suppresses epithelial-mesenchymal transition and stemness markers via activating PI3K/AKT signaling pathway in osteosarcoma cells. Pathol Res Pract (2019) 215:152412. doi:10.1016/j.prp.2019.04.004

26. Wang SH, Baker JR. “Apoptosis in thyroid cancer,” in Thyroid Cancer (Second Edition): A Comprehensive Guide to Clinical Management (Humana Press), 55–61. doi:10.1007/978-1-59259-995-0_6

27. Yang H-L, Pan J-X, Sun L, Yeung S-CJ. p21 Waf-1 (Cip-1) enhances apoptosis induced by manumycin and paclitaxel in anaplastic thyroid cancer cells. J Clin Endocrinol Metab (2003) 88:763–772. doi:10.1210/jc.2002-020992

28. Wang SH, Phelps E, Utsugi S, Baker JRJ. Susceptibility of thyroid cancer cells to 7-hydroxystaurosporine-induced apoptosis correlates with Bcl-2 protein level. Thyroid (2001) 11:725–731. doi:10.1089/10507250152484556

29. Rinner B, Siegl V, Purstner P, Efferth T, Brem B, Greger H, Pfragner R. Activity of novel plant extracts against medullary thyroid carcinoma cells. Anticancer Res (2004) 24:495–500.

30. Wei L, Jin Z, Yang S, Xu Y, Zhu Y, Ji Y. TCGA-assembler 2: software pipeline for retrieval and processing of TCGA/CPTAC data. Bioinformatics (2018) 34:1615–1617. doi:10.1093/bioinformatics/btx812

31. Sanchez-Vega F, Mina M, Armenia J, Chatila WK, Luna A, La KC, Dimitriadoy S, Liu DL, Kantheti HS, Saghafinia S, et al. Oncogenic Signaling Pathways in The Cancer Genome Atlas. Cell (2018) 173:321–337 e10. doi:10.1016/j.cell.2018.03.035

32. van der Net JB, Janssens AC, Defesche JC, Kastelein JJ, Sijbrands EJ, Steyerberg EW. Usefulness of genetic polymorphisms and conventional risk factors to predict coronary heart disease in patients with familial hypercholesterolemia. Am J Cardiol (2009) 103:375–380. doi:10.1016/j.amjcard.2008.09.093

33. Dyrskjot L, Reinert T, Algaba F, Christensen E, Nieboer D, Hermann GG, Mogensen K, Beukers W, Marquez M, Segersten U, et al. Prognostic Impact of a 12-gene Progression Score in Non-muscle-invasive Bladder Cancer: A Prospective Multicentre Validation Study. Eur Urol (2017) 72:461–469. doi:10.1016/j.eururo.2017.05.040

34. Chaudhary K, Poirion OB, Lu L, Garmire LX. Deep Learning-Based Multi-Omics Integration Robustly Predicts Survival in Liver Cancer. Clin Cancer Res (2018) 24:1248–1259. doi:10.1158/1078-0432.CCR-17-0853

35. Pedregosa F, Varoquaux G, Gramfort A, Michel V, Thirion B, Grisel O, Blondel M, Prettenhofer P, Weiss R, Dubourg V, et al. Scikit-Learn: Machine Learning in Python. J Mach Learn Res (2011) 12:2825–2830.

36. Singh H, Kumar R, Singh S, Chaudhary K, Gautam A, Raghava GP. Prediction of anticancer molecules using hybrid model developed on molecules screened against NCI-60 cancer cell lines. BMC Cancer (2016) 16:77. doi:10.1186/s12885-016-2082-y

37. Nagpal G, Usmani SS, Dhanda SK, Kaur H, Singh S, Sharma M, Raghava GPS. Computer-aided designing of immunosuppressive peptides based on IL-10 inducing potential. Sci Rep (2017) 7:42851. doi:10.1038/srep42851

38. Lathwal A, Arora C, Raghava GPS. Prediction of risk scores for colorectal cancer patients from the concentration of proteins involved in mitochondrial apoptotic pathway. PLoS One (2019) 14:e0217527. doi:10.1371/journal.pone.0217527

39. Kaur D, Arora C, Raghava GPS. A Hybrid Model for Predicting Pattern Recognition Receptors Using Evolutionary Information. Front Immunol (2020) 11:71. doi:10.3389/fimmu.2020.00071

40. Dhall A, Patiyal S, Kaur H, Bhalla S, Arora C, Raghava GPS. Computing Skin Cutaneous Melanoma Outcome From the HLA-Alleles and Clinical Characteristics. Front Genet (2020) 11:221. Available at: https://www.frontiersin.org/article/10.3389/fgene.2020.00221

41. Li P, Ren H, Zhang Y, Zhou Z. Fifteen-gene expression based model predicts the survival of clear cell renal cell carcinoma. Med (2018) 97:e11839. doi:10.1097/MD.0000000000011839

42. Wang Y, Ren F, Chen P, Liu S, Song Z, Ma X. Identification of a six-gene signature with prognostic value for patients with endometrial carcinoma. Cancer Med (2018) 7:5632–5642. doi:10.1002/cam4.1806

43. Tang Z, Li C, Kang B, Gao G, Li C, Zhang Z. GEPIA: a web server for cancer and normal gene expression profiling and interactive analyses. Nucleic Acids Res (2017) 45:W98–W102. doi:10.1093/nar/gkx247

44. Stelzer G, Rosen N, Plaschkes I, Zimmerman S, Twik M, Fishilevich S, Stein TI, Nudel R, Lieder I, Mazor Y, et al. The GeneCards Suite: From Gene Data Mining to Disease Genome Sequence Analyses. Curr Protoc Bioinforma (2016) 54:1.30.1–1.30.33. doi:10.1002/cpbi.5

45. Abbott KL, Nyre ET, Abrahante J, Ho Y-Y, Isaksson Vogel R, Starr TK. The Candidate Cancer Gene Database: a database of cancer driver genes from forward genetic screens in mice. Nucleic Acids Res (2015) 43:D844–8. doi:10.1093/nar/gku770

46. Kazaure HS, Roman SA, Sosa JA. The impact of age on thyroid cancer staging. Curr Opin Endocrinol Diabetes Obes (2018) 25:330–334. doi:10.1097/MED.0000000000000430

47. Tuttle RM, Haugen B, Perrier ND. Updated American Joint Committee on Cancer/Tumor-Node-Metastasis Staging System for Differentiated and Anaplastic Thyroid Cancer (Eighth Edition): What Changed and Why? Thyroid (2017) 27:751–756. doi:10.1089/thy.2017.0102

48. Kauffmann RM, Hamner JB, Ituarte PHG, Yim JH. Age greater than 60 years portends a worse prognosis in patients with papillary thyroid cancer: should there be three age categories for staging? BMC Cancer (2018) 18:316. doi:10.1186/s12885-018-4181-4

49. Zhao N, Guo M, Wang K, Zhang C, Liu X. Identification of Pan-Cancer Prognostic Biomarkers Through Integration of Multi-Omics Data. Front Bioeng Biotechnol (2020) 8:268. doi:10.3389/fbioe.2020.00268

50. Shen Y, Dong S, Liu J, Zhang L, Zhang J, Zhou H, Dong W. Identification of Potential Biomarkers for Thyroid Cancer Using Bioinformatics Strategy: A Study Based on GEO Datasets. Biomed Res Int (2020) 2020:9710421. doi:10.1155/2020/9710421

51. Musa A, Ghoraie LS, Zhang S-D, Glazko G, Yli-Harja O, Dehmer M, Haibe-Kains B, Emmert-Streib F. A review of connectivity map and computational approaches in pharmacogenomics. Brief Bioinform (2018) 19:506–523. doi:10.1093/bib/bbw112

52. Lamb J, Crawford ED, Peck D, Modell JW, Blat IC, Wrobel MJ, Lerner J, Brunet J-P, Subramanian A, Ross KN, et al. The Connectivity Map: using gene-expression signatures to connect small molecules, genes, and disease. Science (2006) 313:1929–1935. doi:10.1126/science.1132939

53. Shinwari Z, Manogaran PS, Alrokayan SA, Al-Hussein KA, Aboussekhra A. Vincristine and lomustine induce apoptosis and p21(WAF1) up-regulation in medulloblastoma and normal human epithelial and fibroblast cells. J Neurooncol (2008) 87:123–132. doi:10.1007/s11060-007-9502-4

54. Buss JL, Torti FM, Torti S V. The role of iron chelation in cancer therapy. Curr Med Chem (2003) 10:1021–1034. doi:10.2174/0929867033457638

55. Marques O, da Silva BM, Porto G, Lopes C. Iron homeostasis in breast cancer. Cancer Lett (2014) 347:1–14. doi:10.1016/j.canlet.2014.01.029

56. Yang Y, Xu Y, Su A, Yang D, Zhang X. Effects of Deferoxamine on Leukemia In Vitro and Its Related Mechanism. Med Sci Monit (2018) 24:6735–6741. doi:10.12659/MSM.910325

57. Bajbouj K, Shafarin J, Hamad M. High-Dose Deferoxamine Treatment Disrupts Intracellular Iron Homeostasis, Reduces Growth, and Induces Apoptosis in Metastatic and Nonmetastatic Breast Cancer Cell Lines. Technol Cancer Res Treat (2018) 17:1533033818764470. doi:10.1177/1533033818764470

58. Petrella A, Festa M, Ercolino SF, Zerilli M, Stassi G, Solito E, Parente L. Annexin-1 downregulation in thyroid cancer correlates to the degree of tumor differentiation. Cancer Biol Ther (2006) 5:643–647. doi:10.4161/cbt.5.6.2700

59. Fuzio P, Napoli A, Ciampolillo A, Lattarulo S, Pezzolla A, Nuzziello N, Liuni S, Giorgino F, Maiorano E, Perlino E. Clusterin transcript variants expression in thyroid tumor: a potential marker of malignancy? BMC Cancer (2015) 15:349. doi:10.1186/s12885-015-1348-0

60. Chang Y-S, Chang C-C, Huang H-Y, Lin C-Y, Yeh K-T, Chang J-G. Detection of Molecular Alterations in Taiwanese Patients with Medullary Thyroid Cancer Using Whole-Exome Sequencing. Endocr Pathol (2018) 29:324–331. doi:10.1007/s12022-018-9543-6

61. Lian M, Cao H, Baranova A, Kural KC, Hou L, He S, Shao Q, Fang J. Aging-associated genes TNFRSF12A and CHI3L1 contribute to thyroid cancer: An evidence for the involvement of hypoxia as a driver. Oncol Lett (2020) 19:3634–3642. doi:10.3892/ol.2020.11530

62. Qiu J, Zhang W, Zang C, Liu X, Liu F, Ge R, Sun Y, Xia Q. Identification of key genes and miRNAs markers of papillary thyroid cancer. Biol Res (2018) 51:45. doi:10.1186/s40659-018-0188-1

63. McCann JC, Ames BN. Adaptive dysfunction of selenoproteins from the perspective of the triage theory: why modest selenium deficiency may increase risk of diseases of aging. FASEB J (2011) 25:1793–1814. doi:10.1096/fj.11-180885

64. Zarkesh M, Zadeh-Vakili A, Azizi F, Fanaei SA, Foroughi F, Hedayati M. The Association of BRAF V600E Mutation With Tissue Inhibitor of Metalloproteinase-3 Expression and Clinicopathological Features in Papillary Thyroid Cancer. Int J Endocrinol Metab (2018) 16:e56120. doi:10.5812/ijem.56120

65. Dong T, Zhang Z, Zhou W, Zhou X, Geng C, Chang LK, Tian X, Liu S. WNT10A/betacatenin pathway in tumorigenesis of papillary thyroid carcinoma. Oncol Rep (2017) 38:1287–1294. doi:10.3892/or.2017.5777

66. Gugnoni M, Sancisi V, Gandolfi G, Manzotti G, Ragazzi M, Giordano D, Tamagnini I, Tigano M, Frasoldati A, Piana S, et al. Cadherin-6 promotes EMT and cancer metastasis by restraining autophagy. Oncogene (2017) 36:667–677. doi:10.1038/onc.2016.237

