## Supplementary S2 for "Prognostic Biomarkers for Predicting Papillary Thyroid Carcinoma Patients at High Risk Using Nine Genes of Apoptotic Pathway"

**Supplementary File S2**

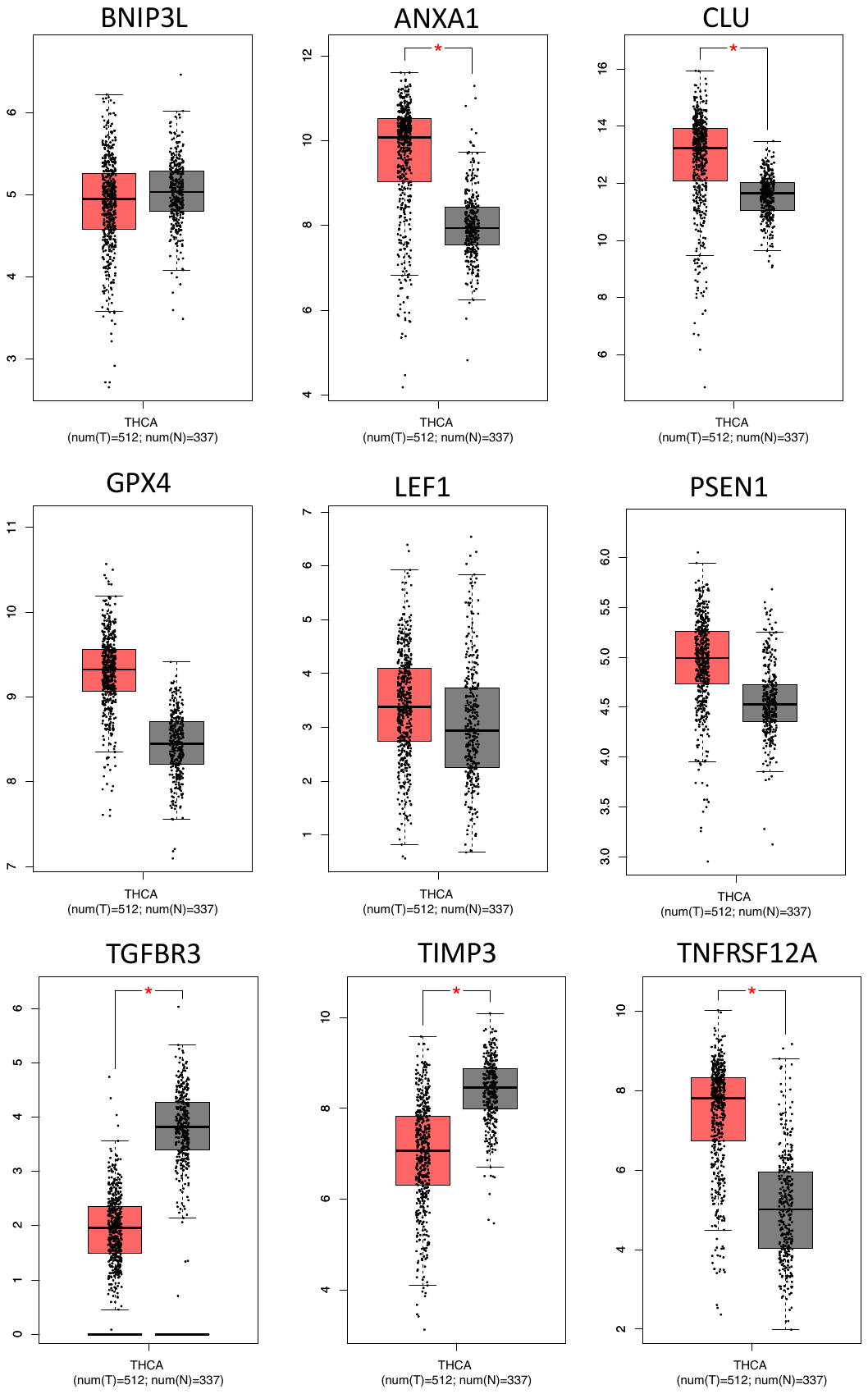

**Figure A** Boxplots representing the differential gene expression between normal and tumour samples on a log scale. GEPIA webserver was used to plot these by using TCGA THCA dataset. T: Tumour in red, N: Normal (TCGA,GTEX) in grey.

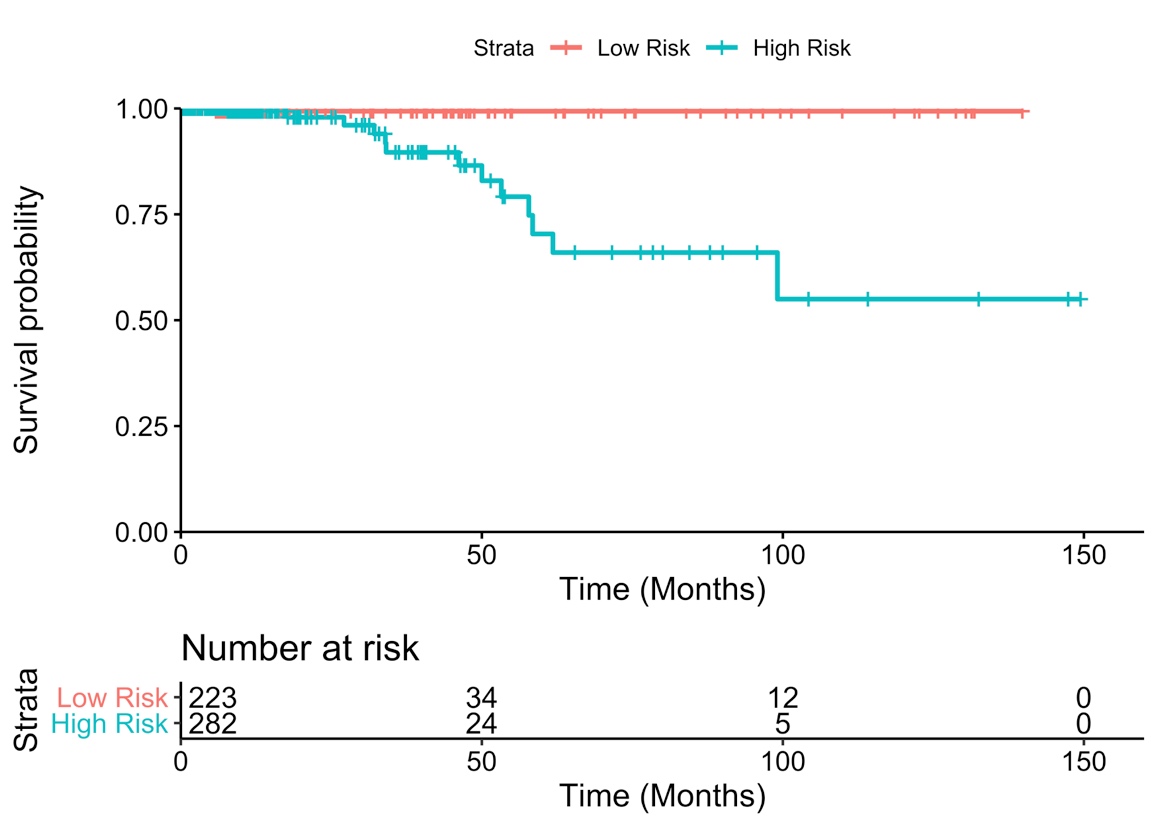

**Figure B** Risk stratification of PTC patients using prognostic index (PI) model. Patients with PI> -3.29x10^5^ (estimated using cutp() package in R) were found to be at higher risk with HR=17.55, p=5.88x10^-3^, C=0.65, %95CI 2.29-134.72 and logrank-p=6.73x10^-5^.

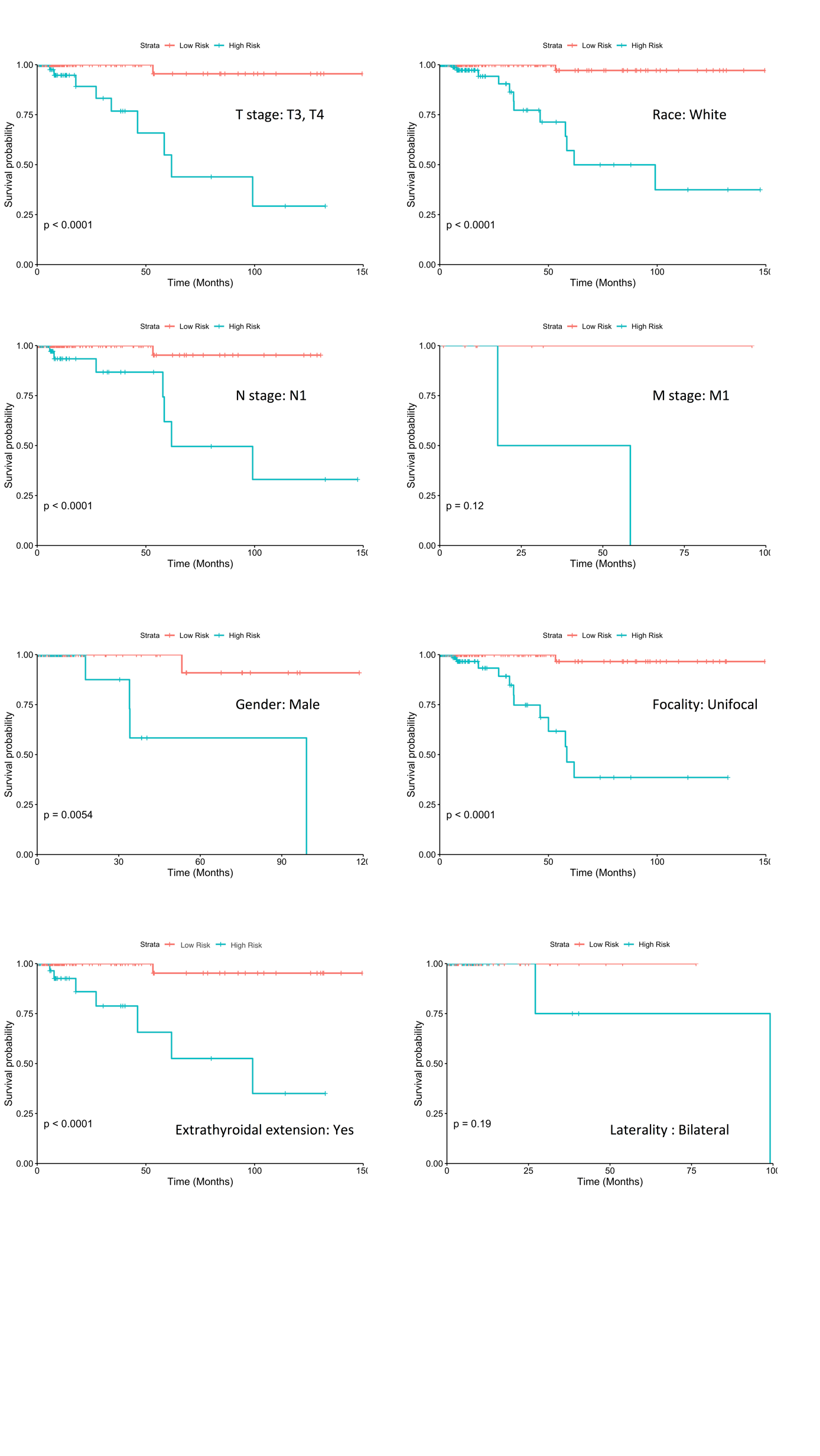

**Figure C** Sub-stratification of clinico-pathological high risk groups by 9-gene voting model. Logrank p values show significant segregation between survival curves.
